# Cost-free lifespan extension via optimisation of gene expression in adulthood supports the developmental theory of ageing

**DOI:** 10.1101/739490

**Authors:** Martin I. Lind, Hanne Carlsson, Edward Ivimey-Cook, Alexei A. Maklakov

**Affiliations:** Animal Ecology, Department of Ecology and Genetics, Uppsala University, Uppsala, SE-75236, Sweden; School of Biological Sciences, University of East Anglia, Norwich Research Park, Norwich NR4 7TJ, UK

**Keywords:** ageing, antagonistic pleiotropy, age-specific fitness, IIS signalling, TOR signalling, protein synthesis, mitochondrial respiration, life-history evolution, senescence

## Abstract

Classic theory upholds that energy trade-offs between reproduction and somatic maintenance underpin the evolution of ageing and lifespan. In contrast, the developmental theory of ageing (DTA) suggests that organismal senescence is caused by dysregulated gene expression in adulthood due to decline in selection gradients with age. The DTA predicts that age-specific optimisation of gene expression can improve survival without fitness costs. Here we investigated consequences for survival, reproduction, egg size and fitness of early-life, adulthood and post-reproductive onset of RNAi knockdown of five well-described “longevity” genes involved in key biological processes in *Caenorhabditis elegans* nematodes: nutrient-sensing signalling via insulin/IGF-1 (*age-1*) and target-of-rapamycin (*raga-1*) pathways, global protein synthesis (*ifg-1*), global protein synthesis in somatic cells (*ife-2*) and mitochondrial respiration (*nuo-6*). Downregulation of these genes in adulthood and/or during post-reproductive period improves survival, while there was little evidence for a link between impaired reproduction and extended lifespan. Our findings demonstrate that hyper-function of diverse physiological processes after sexual maturation is detrimental for survival. Therefore, optimisation of gene expression in adult organisms can ameliorate ageing and increase fitness.

## Results and Discussion

The force of natural selection is maximised during pre-reproductive development but declines after sexual maturation with advancing age [1-4]. Therefore, antagonistically pleiotropic alleles that have positive fitness effects early in life but negative fitness effects late in life can be selected for and lead to the evolution of ageing [1]. While the antagonistic pleiotropy (AP) theory is widely accepted, the proximate routes that lead to ageing are poorly understood. The dominant paradigm, the “disposable soma” theory of ageing (DST), postulates that ageing and longevity evolve as a result of optimised energy allocation between somatic maintenance and reproduction with the aim of maximising the reproductive output [5-7]. This theory predicts that increased investment in soma will increase survival at the cost of reduced reproduction, and vice versa. Indeed, there is corroborating evidence from laboratory [reviewed in 8] and field [reviewed in 9] studies suggesting that there is a link between increased reproduction and reduced lifespan. Nevertheless, the predominance of this theory has been increasingly challenged in recent years, both empirically [10-14] and theoretically [15-17]. Several studies in different model organisms have suggested that increased longevity and reduced reproduction can be uncoupled, thereby questioning the key role of energy allocation trade-offs in ageing. In contrast, the emerging developmental theory of ageing (DTA) maintains that the decline in selection gradients with age results in suboptimal regulation of gene expression in late adulthood leading to cellular and organismal senescence. The DTA upholds that gene expression is optimised for development and early-life reproduction and, therefore, if gene expression is instead optimised across the whole life course of the organism, survival can be improved without detrimental effects on other fitness-related traits, such as propagule size and offspring number.

Here we tested this prediction directly by modifying age-specific expression of five well-described “longevity” genes in *Caenorhabditis elegans* nematode worms that play key roles in different physiological processes: nutrient-sensing signalling via insulin/IGF-1 (*age-1*) [18, 19] and target-of-rapamycin (*raga-1*) [20, 21] pathways, global protein synthesis (*ifg-1*) [22], global protein synthesis in somatic cells (*ife-2*) [23, 24] and mitochondrial respiration (*nuo-6*) [25]. The *age-1* gene encodes the phosphatidylinositol 3-kinase (PI3K) catalytic subunit homologue, which is involved in kinase-phosphorylation cascade that downregulates the DAF16/FOXO transcription factor [19]. Loss-of-function mutations in *age-1* increase lifespan [18, 19] but reduce early-life reproduction and fitness [26-28]. The *raga-1* encodes *C. elegans* orthologue of GTPase RagA, which is the amino-acid sensing activator of the target-of-rapamycin complex 1 (TORC1) signal transduction pathway [20] that governs cell growth and shapes lifespan [29]. Loss-of-function *raga-1* mutants have longer lifespan and slower behavioural decline with age [30]. The *ife-2* encodes a eukaryotic translation initiation factor eIF4E, which is a regulator of protein synthesis and is most abundant in the somatic cells in *C. elegans*. Under standard temperature (20°C), disruption of *ife-2* via mutation or lifelong RNA interference (RNAi) increases survival and without negative effects on brood size [24, 31], and it is suggested that the lifespan extension is conferred specifically via reduction of protein synthesis in the soma [32]. The *nuo-6* encodes mitochondrial subunit of complex I in the mitochondrial respiratory chain, and lifelong *nuo-6* RNAi reduces growth and fertility but increases longevity [25].

Our approach was to use age-specific RNAi to downregulate the expression of these genes starting at three different stages across the life course of *C. elegans*: i) newly laid egg (lifelong treatment), ii) sexual maturity (adulthood treatment) and iii) the end of self-fertilized reproduction (post-reproductive treatment). This approach allowed us to assess the fitness consequences of lifelong and adulthood-only downregulation of the target genes, as well as the effects of post-reproductive downregulation on survival. The latter effect is particularly interesting in this regard, because it allows us to test whether age-specific optimisation of gene function can extend lifespan in the absence of the energy cost of reproduction. If post-reproductive downregulation of gene expression can indeed increase survival, it would be a proof-of-principle that gene expression in late-life is not optimised for long life. Nevertheless, an even more crucial test was whether we can modify different physiological functions in adulthood to improve survival without hampering key life-history traits. We investigated the age-specific RNAi effects on survival, age-specific reproduction and egg size (as a measure of parental investment into offspring and a proxy for offspring quality); we then used these data to determine lifetime reproductive success (LRS) and rate-sensitive individual fitness (λ_ind_).

Timing of RNAi treatment had profound effects on survival, age-specific and lifetime reproduction, egg size and fitness (Fig.1 and 2, Fig. S1, Tables 1 and 2). Nevertheless, there was little evidence for a link between increased lifespan and reduced fitness. Downregulation of *age-1* across all life stages improved longevity but the effect became progressively weaker with increasing age of onset of the RNAi treatment; however, we found no indication that *age-1* RNAi negatively affected lifetime reproductive success (LRS), egg size or individual fitness (λ_ind_) (Fig. 1 and 2, Fig. S1, Table 1). Interestingly, the effect of TORC1 downregulation via *raga-1* RNAi on traits was quite different: lifelong RNAi did not have any positive effect, while adulthood-only and post-reproductive treatments slightly improved survival but did not affect LRS, egg size or individual fitness (Fig.1 and 2, Fig. S1, Table 1). These results suggest that the two major nutrient-sensing molecular signalling pathways, IIS and TOR, have very different effects on vital life-history traits.

**Table 1.** The overall effect of down-regulating *age-1, raga-1, nuo-6, ifg-1* or *ife-2* on lifespan, fitness (λ_ind_), lifetime reproductive success (LRS) and egg size.

| Gene | Factor | **Lifespan** |  |  | **Fitness ($\lambda_{ind}$)** |  |  | **LRS** |  |  | **Egg size** |  |  |
| --- | --- | --- | --- | --- | --- | --- | --- | --- | --- | --- | --- | --- | --- |
| | | $\chi^2$ | d.f. | p | $\chi^2$ | d.f. | p | $\chi^2$ | d.f. | p | $\chi^2$ | d.f. | p |
| *age-1* | Treatment | 137.97 | 3 | **<0.001** | 4.838 | 2 | 0.089 | 1.484 | 2 | 0.476 | 0.26 | 2 | 0.878 |
| *raga-1* | Treatment | 45.681 | 3 | **<0.001** | 0.986 | 2 | 0.611 | 3.024 | 2 | 0.221 | 3.30 | 2 | 0.192 |
| *nuo-6* | Treatment | 206.72 | 3 | **<0.001** | 66.352 | 2 | **<0.001** | 93.681 | 2 | **<0.001** | 9.40 | 2 | **0.009** |
| *ifg-1* | Treatment | 108.43 | 3 | **<0.001** | 541.13 | 2 | **<0.001** | 1271.300 | 2 | **<0.001** | 151.65 | 2 | **<0.001** |
| *ife-2* | Treatment | 110.89 | 3 | **<0.001** | 5.485 | 2 | 0.064 | 0.707 | 2 | 0.702 | 6.45 | 2 | **0.040** |

**Table 2.** Age-specific reproduction. The effect of RNAi-treatment, Age and Age^2^ on age-specific reproduction for each of the five genes. All models were fitted using a Conway-Maxwell-Poisson (CMP) distribution (models with CMP distribution had lowest AIC for all genes, see Table S7). It was not possible to model age-specific reproduction for *ifg-1*, since most treatment-levels and ages lacked reproduction (see Fitness and LRS instead).

| Gene | Factor | *Age-specific reproduction* |  |  |
| --- | --- | --- | --- | --- |
| | | $\chi^2$ | d.f. | p |
| *age-1* | Treatment | 0.704 | 2 | 0.704 |
|  | Age | 1094.192 | 1 | **<0.001** |
|  | Age<sup>2</sup> | 1115.490 | 1 | **<0.001** |
|  | Treatment × Age | 5.826 | 2 | 0.054 |
|  | Treatment × Age<sup>2</sup> | 5.174 | 2 | 0.075 |
| *raga-1* | Treatment | 1.536 | 2 | 0.464 |
|  | Age | 1202.446 | 1 | **<0.001** |
|  | Age<sup>2</sup> | 1248.191 | 1 | **<0.001** |
|  | Treatment × Age | 11.888 | 2 | **0.003** |
|  | Treatment × Age<sup>2</sup> | 11.186 | 2 | **0.004** |
| *nuo-6* | Treatment | 79.657 | 2 | **<0.001** |
|  | Age | 522.752 | 1 | **<0.001** |
|  | Age<sup>2</sup> | 589.162 | 1 | **<0.001** |
|  | Treatment × Age | 18.537 | 2 | **<0.001** |
|  | Treatment × Age<sup>2</sup> | 17.870 | 2 | **<0.001** |
| *ifg-1* | *Almost complete cross-separation, not possible to model* |  |  |  |
| *ife-2* | Treatment | 0.408 | 2 | 0.815 |
|  | Age | 1439.479 | 1 | **<0.001** |
|  | Age<sup>2</sup> | 1487.427 | 1 | **<0.001** |
|  | Treatment × Age | 16.522 | 2 | **<0.001** |
|  | Treatment × Age<sup>2</sup> | 23.244 | 2 | **<0.001** |

**Figure 1.**
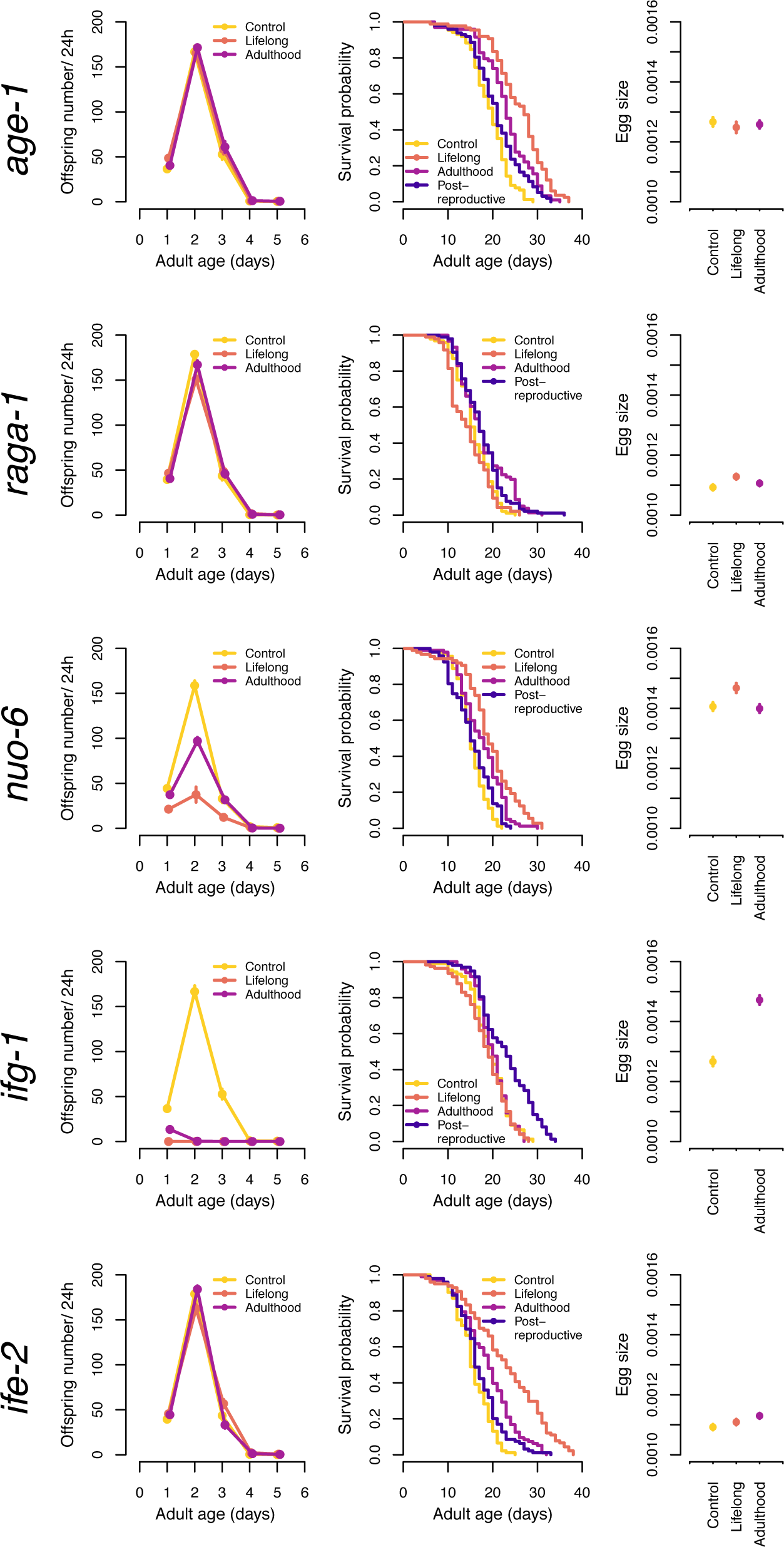
Age-specific reproduction, lifespan and egg size, presented for each gene, and separated by treatment group. Colours indicate control (yellow), lifelong RNAi treatment (orange), RNAi during adulthood only (purple) and post-reproductive RNAi (blue). For age-specific reproduction and egg size, symbols represent mean ± SE.

**Figure 2.**
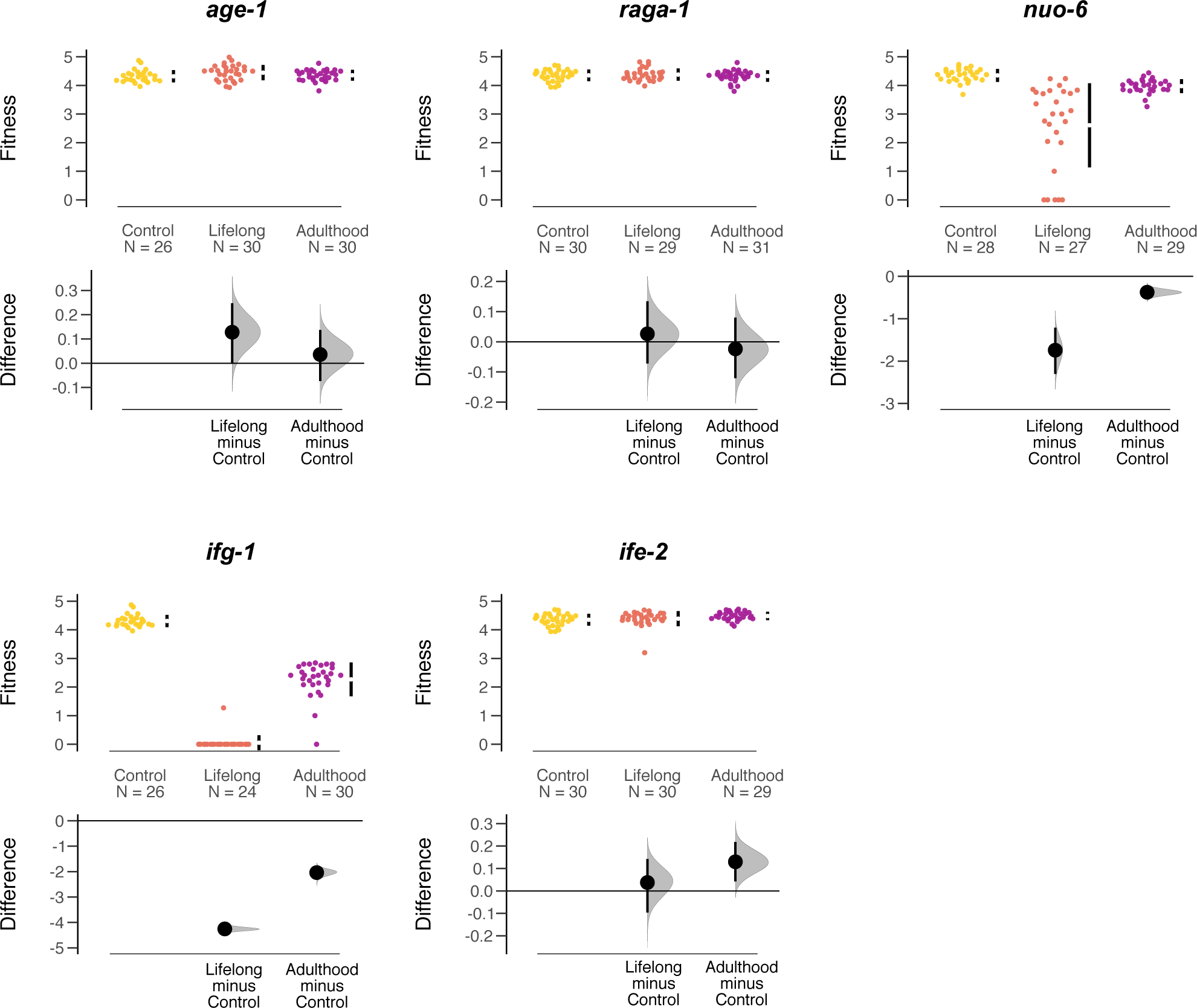
Individual fitness (λind) calculated from age-specific fecundity data, separated by gene and treatment group: control (yellow), lifelong RNAi treatment (orange) and RNAi during adulthood only (purple). Top panels show raw data, with the mean ± 95%CI indicated by black bars at each group. Bottom panels show estimation plots, where RNAi treatments are compared to the control, with a graded sampling distribution of bootstrapped values and the bootstrapped 95% CI.

Age-specific downregulation of *nuo-6* showed a perfect negative correlation between survival and reproduction. Similar to the results with *age-1, nuo-6* RNAi increased survival and the effect became weaker with increased age of onset of RNAi treatment (Fig.1, Table 2). Contrary to effect of *age-1* RNAi, however, improved survival was mirrored by negative effects on LRS and fitness, while egg size was improved in the lifelong treatment (Fig.1 and 2, Fig. S1, Tables 1 and 2).

Downregulation of *ifg-1* predictably abolished reproduction in the lifelong treatment, and severely reduced it when started in adulthood, while the few eggs produced in adulthood-only treatment were quite large (Fig. 1, Fig. S1, Table 1). Interestingly, there was no effect of reduced reproduction on survival (Fig. 1, Table 2). Perhaps even more remarkable was the positive effect of post-reproductive *ifg-1* RNAi on survival (Fig.1, Table 2). These results support the notion that superfluous protein synthesis in late-life reduces longevity in *C. elegans*.

Age-specific downregulation of *ife-2* increased survival across all treatments with the effect becoming progressively weaker with the later age of RNAi onset (Fig.1, Table 1), similar to the results with *age-1* and *nuo-6*. Interestingly, there were no negative effects on LRS (Fig. S1, Table 1), while adulthood-only RNAi actually increased egg size (Fig. 1, Tables 1 and 2). Thus, adulthood-only *ife-2* RNAi simultaneously improved survival and investment in offspring; moreover, within-model contrasts suggest adulthood-only *ife-2* RNAi had higher individual fitness than control animals (Table S4), although there was no overall significant effect across all three treatments (Table 1). Nevertheless, bootstrapping analyses which does not depend on a specified distribution of the data suggests that adulthood-only *ife-2* RNAi does increase rate-sensitive fitness (Fig. 2).

The dominant DST paradigm proposes that improved somatic maintenance necessitates increased energy allocation, which will lead to reduced investment in growth and reproduction. Contrary to this, the DTA maintains that survival can be improved by optimising age-specific gene expression without reproduction costs because gene expression is predicted to be optimised for development and early-life reproduction. The corollary of this argument that optimising gene expression during adulthood can even increase Darwinian fitness. The force of natural selection declines with age, and does so very rapidly in small fast-reproducing organisms such as *C. elegans* [33]. This means that even very small positive effects on vital life-history traits early in life can be beneficial for Darwinian fitness despite large fitness costs late in life [2, 33]. This also implies that natural selection on regulating gene expression in late-life is very weak in *C. elegans* and there is scope for experimental optimisation of age-specific gene expression.

We found that most of the “longevity” genes that we tested showed poor correlation between the age-specific gene expression effects in survival, reproduction, egg size and individual fitness (Fig. 1 and 2, Table 1). Only one of these genes – *nuo-6* – showed the pattern of the negative correlation between increase in survival and reduced reproduction and fitness that is predicted under the DST. We note that such correlation does not imply causation even in this one case, and it is possible that upregulation of stress resistance pathways in *nuo*-6 does not depend on energy reallocation from reduced egg laying. These results support the previous work showing that rates of ageing are affected by mitochondrial function during development [34]. Nevertheless, our result also showed that adulthood-only reduction in mitochondrial respiration also can extend lifespan. We also note that the lack of negative effects of RNAi on LRS or fitness in other genes was not caused by lack of power to detect negative effects, because the treatment means of these non-significant effects were actually positive for most comparisons (Figure 2, S1).

Remarkably, adulthood-only RNAi knockdown of *ife-2* improved survival and egg size. These results suggest that superfluous protein synthesis in the somatic cells of adult worms promotes cellular senescence and reduces Darwinian fitness through the effects on both parents and their offspring through egg size.

More generally, we showed that adulthood-only, or even post-reproductive downregulation of important physiological functions, such as IIS nutrient-sensing signalling, global protein synthesis in all tissues and in the somatic cells, and mitochondrial respiration can improve survival without negative fitness effects. The results that we obtained in age-specific *age-1* and *ife-2* RNAi experiments suggest that adulthood-only knockdowns can increase fitness because they improve survival without negative effects on reproduction, and even a positive effect on egg size and individual fitness in the case of *ife-2*. Perhaps particularly intriguing is the fact that in four out of five cases, lifespan extension could be achieved via post-reproductive onset of RNAi treatment. While post-reproductive worms do not affect the allelic frequencies in the next generation, as there is no post-hatching parental care in this system, this latter result nevertheless suggests that late-life hyperfunction of certain genes contributes to earlier death.

Overall, the results of this study are consistent with the hypothesis that selection optimises gene expression in early life, while post-maturation expression can be optimised further, as predicted by the developmental theory of ageing. While the lack of selection on gene expression during post-reproductive period is rather straightforward, one can question why is selection so weak during the reproductive period of *C. elegans* life cycle. The answer likely lies in the biology of this species, which is characterised by a very rapid and strong (orders of magnitude) age-specific decline in selection gradients [33]. Indeed, the selection gradients on fecundity decline nearly exponentially with age in the laboratory [33], and this decline is likely further exacerbated in nature where the food resources are ephemeral. Therefore, small differences in fitness of two-day old worms may be largely invisible to selection. However, while the decline in selection gradients with age is particularly strong in *C. elegans*, reduced force of selection with advancing age is a general pattern across organisms.

## Conclusions

Our findings support the hypothesis that gene expression is optimised for development and early-life reproduction across a broad range of physiological processes. Consequently, gene expression in adulthood can be optimised further to improve survival and, potentially, fitness. Only *ife-2* adulthood-only RNAi animals had simultaneously increased survival and egg size; however, longevity usually correlates with increased resistance to different ecologically relevant stressors, such as temperature, light and pathogens [35, 36], so it is likely that improved survival could contribute positively to fitness under challenging conditions in nature. These results of course do not preclude the possibility that other types of trade-offs contribute to ageing in *C. elegans*. One important aspect to consider is phenotypic plasticity and how these animals would perform in different contexts. Future work should focus on studying fitness consequences of age-specific gene expression optimisation across a broad range of ecologically relevant environments.

Notwithstanding the results of such future studies, these findings here strongly support the theoretical conjecture that non-energy-based trade-offs between gene effects on fitness across the life course play a key role in the evolution and expression of ageing and longevity.

## Acknowledgements

This work has been supported by BBSRC BB/R017387/1 and ERC Consolidator Grant GermlineAgeingSoma to AAM and Swedish Research Council grant 2016-05195 (Vetenskapsrådet) to MIL.

## Methods

### Strains

*Caenorhabditis elegans* nematodes of strain Bristol N2 wild-type, obtained from Caenorhabditis Genetics Center, were used in all assays. Populations were recovered from frozen and bleached before the start of the experiment. Standard NGM agar plates [1] were used to grow the nematode populations and antibiotics (100 μg/ml Ampicillin and 100 μg/ml Streptomycin) and a fungicide (10 μg/ml Nystatin) was added to the agar to avoid infections [2]. Up until the start of the experiment nematode populations were fed antibiotic resistant *Echerichia coli* OP50-1 (pUC4K), gifted by J. Ewbank at the Centre d’Immunologie de Marseille-Luminy, France. During recovery from freezing and throughout the experiment the worms were retained in climate chambers maintaining 20°C and 60% relative humidity.

In order to induce RNAi knockdown, the nematodes were fed *E. coli* of the strain HT115 (DE3) containing a Timmons and Fire feeding vector L4440 modified to express the dsRNA of the gene of interest. The genes targeted by RNAi were age-1(B0334.8), raga-1 (T24F1.1), nuo-6 (W01A8.4), ife-2 (R04A9.4) and ifg-1 (M110.4). These strains were provided by Source Bioscience and Julie Ahringer. In addition to the RNAi knockdown bacteria, a control strain of HT115 was used, carrying an empty L4440 feeding vector [3].

Cultures of the RNAi clones were grown in LB medium supplemented with Ampicillin (50 μg/ml) before seeded onto 35mm standard NGM plates with the addition of IPTG (1 mM) and Ampicillin (50 μg/ml) as recommended by Kamath et al. [4]. After seeding, the bacteria was allowed to grow over night in 20°C to induce expression of RNAi, before worms were placed on the plates.

### Experimental set-up

Age synchronized eggs were collected from OP50-1 (pUC4K) fed unmated hermaphrodites on adult day 2. These eggs were placed either on RNAi seeded plates and maintained on RNAi throughout life (*lifelong-exposure*), on empty vector from egg to late L4 stage and then onto RNAi (*adulthood-exposure*), on empty vector from egg to day 6 of adulthood and then on RNAi (*post-reproductive exposure*) or maintained throughout life on empty vector plates (*control*). For every gene knockdown, assays were performed in two blocks. The scoring was achieved by a blinded observer, with agar plates of the different treatments handled in a randomized order.

### Reproductive assays

To gather daily reproductive output per worm, unmated hermaphrodites were reared on individual plates from late L4 stage until reproduction ceased. The worms were moved onto new plates every 24h. Eggs laid on the plates were allowed to hatch and develop during two days, when the total amount of worms on the plates were counted. For each block, 15 replicate worms were set up for each gene and treatment combination, giving 30 worms per gene, except for the second block of ife-2 and raga-1, where more worms were set up in order to compensate for lost worms in the first block.

### Lifespan assays

Lifespan assays were performed on unmated hermaphrodites from larval stage L4 until death. Worms were set up in groups of ten and transferred onto fresh plates daily while scoring survival. Death was defined as the absence of movement in response to touch. For each block, 50 replicate worms were set up for each gene and treatment combination, giving 100 worms per gene, except for the second block of ife-2 and raga-1, where more worms were set up in order to compensate for lost worms in the first block.

### Egg size measurement

Egg photos were taken from the same plates as used in the lifespan assays. At adult day 2 the 10 lifespan worms were allowed to lay eggs for 2 hours on a fresh agar plate, after which photos of 10 eggs were taken per plate of 10 worms. A microscope camera was used to attain the photos which were later analysed in *ImageJ* (**https://imagej.nih.gov/ij/**).

### Statistical analyses

All analyses were performed separately for each gene. Before analysis, we excluded individuals from plates that were severely contaminated by infection. We also removed one infertile control individual (table S1-S2).

All statistical analyses were performed using the statistical software R 3.6.0. Lifespan was analysed using Cox proportional hazard models implemented in the coxme package, with Treatment as a fixed factor and Block and Plate as a random effects. Individuals dying of matricide (internal hatching of eggs) were censored.

Age-specific reproduction was analysed using generalized linear mixed-effect models. We used the first 3 days of reproduction, since reproduction ceased at day 4. We treated Treatment, Age and Age2 as crossed fixed factors, and fitted Block as well as Individual as random effects (to control for repeated measures). Since reproduction data is often overdispersed, we fitted three different model implementations. First, we fitted the models using a Poisson distribution in the lme4 package. Secondly, we also included a subject-level random effect in the model, to control for possible overdispersion. Thirdly, we fitted a model with a Conway-Maxwell-Poisson (CMP) distribution using the glmmTMB package, which numerically estimates the mean and variance separately, and is well suited to deal with overdispersed data [5]. The models where then compared using AIC and the model with lowest AIC selected. Treatment levels without any reproduction were removed in order to run the model (lifelong exposure to ifg-1).

Individual fitness (λind) was calculated from the life-table of age-specific reproduction, by solving the Euler-lotka equation using the lambda function in the popbio package. The life-table was constructed using a development time of two days. We then analysed λind in linear mixed-effect models using the lme4 package, with treatment as a Fixed factor and Block as a random effect.

Lifetime reproductive success (LRS) was scored as the total number of offspring per individual, and was analysed using generalized linear mixed-effect models using Poisson or CMP distribution, as described above for age-specific reproduction. We fitted treatment as a fixed factor and Block as a random effect.

Individual fitness and LRS was also analysed by bootstrapping, using the dabestrR package, and the 95% confidence intervals are graphically presented by the package in figure 2 and S1.

Egg size was analysed in linear mixed-effect models using the lme4 package, with Treatment as a fixed effect and Block and Plate as random effects. Treatments without any eggs produced were not included (lifelong exposure to ifg-1).

