## Supplementary for "Cost-free lifespan extension via optimisation of gene expression in adulthood supports the developmental theory of ageing"

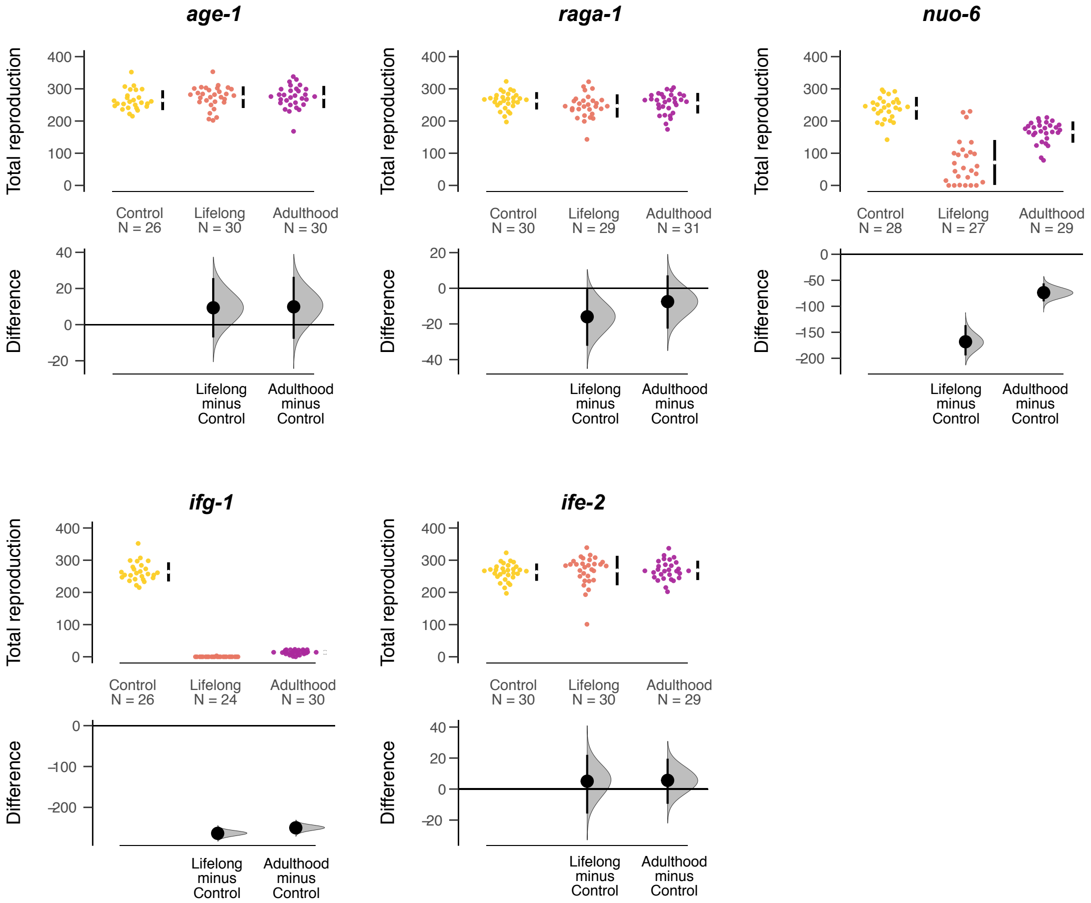


Figure S1. Lifetime reproductive success (LRS) separated by gene and treatment group: control (yellow), lifelong RNAi treatment (orange) and RNAi during adulthood only (purple). Top panels show raw data, with the mean ± 95%CI indicated by black bars at each group. Bottom panels show estimation plots, where RNAi treatments are compared to the control, with a graded sampling distribution of bootstrapped values and the bootstrapped 95% CI.

**Table S1**. The number of individuals of each gene and treatment combination that were excluded from the *lifespan analysis* because of a bacterial infection on the NGM plate. The numbers are a multiply of 10, since the whole plate was always removed.

| Gene | Control | Lifelong | Adulthood | Post-reproductive |
|---|---|---|---|---|
| *age-1* | 10 | 0 | 0 | 0 |
| *raga-1* | 10 | 0 | 0 | 0 |
| *nuo-6* | 0 | 10 | 0 | 0 |
| *ifg-1* | 10 | 0 | 0 | 0 |
| *ife-2* | 10 | 10 | 20 | 10 |

**Table S2**. The number of individuals of each gene and treatment combination that were excluded from the *reproduction analysis* because of a bacterial infection on the NGM plate.

| Gene | Control | Lifelong | Adulthood |
|---|---|---|---|
| *age-1* | 4 | 0 | 0 |
| *raga-1* | 5 | 2 | 2 |
| *nuo-6* | 0 | 0 | 0 |
| *ifg-1* | 0 | 0 | 0 |
| *ife-2* | 5 | 2 | 7 |

**Table S3. Lifespan.** The effect of age-specific down-regulation of each gene on lifespan. Treatment contrast from Cox proportional hazard models, presented for each gene.

| Gene | Treatment contrast | coef | SE | z | p |
|---|---|---|---|---|---|
| *age-1* | Lifelong | -1.76 | 0.191 | -9.19 | **<0.001** |
| | Adulthood | -1.11 | 0.329 | -6.30 | **<0.001** |
| | Post-reproductive | -0.66 | 0.517 | -3.89 | **<0.001** |
| *raga-1* | Lifelong | 0.135 | 0.191 | 0.71 | 0.480 |
| | Adulthood | -0.647 | 0.202 | -3.21 | **0.001** |
| | Post-reproductive | -0.454 | 0.193 | -2.34 | **0.019** |
| *nuo-6* | Lifelong | -1.52 | 0.218 | -5.49 | **<0.001** |
| | Adulthood | -0.92 | 0.263 | -3.51 | **<0.001** |
| | Post-reproductive | -0.06 | 0.258 | -0.23 | 0.820 |
| *ifg-1* | Lifelong | 0.07 | 0.17 | 0.43 | 0.670 |
| | Adulthood | -0.05 | 0.18 | -0.29 | 0.770 |
| | Post-reproductive | -1.16 | 0.20 | -5.86 | **<0.001** |
| *ife-2* | Lifelong | -1.52 | 0.17 | -8.89 | **<0.001** |
| | Adulthood | -0.69 | 0.50 | -4.59 | **<0.001** |
| | Post-reproductive | -0.39 | 0.68 | -2.61 | **0.009** |

**Table S4. Fitness.** The effect of age-specific down-regulation of each gene on fitness ($\lambda_{ind}$). Treatment contrast from mixed-effect models.

| Gene | Treatment contrast | coef | SE | d.f. | t | p |
|---|---|---|---|---|---|---|
| *age-1* | Intercept (Control) | 4.318 | 0.044 | 83 | 97.29 | **<0.001** |
| | Lifelong | 0.128 | 0.061 | 83 | 2.11 | **0.038** |
| | Adulthood | 0.036 | 0.061 | 83 | 0.59 | 0.556 |
| *raga-1* | Intercept (Control) | 4.342 | 0.050 | 2.49 | 87.52 | **<0.001** |
| | Lifelong | 0.033 | 0.053 | 86.4 | 0.62 | 0.539 |
| | Adulthood | -0.019 | 0.052 | 86.3 | -0.36 | 0.717 |
| *nuo-6* | Intercept (Control) | 4.364 | 0.267 | 1.68 | 16.36 | **0.008** |
| | Lifelong | -1.735 | 0.223 | 80.0 | -7.79 | **<0.001** |
| | Adulthood | -0.396 | 0.219 | 80.0 | -1.81 | 0.074 |
| *ifg-1* | Intercept (Control) | 4.148 | 0.120 | 78 | 34.35 | **<0.001** |
| | Lifelong | -4.095 | 0.176 | 78 | -23.26 | **<0.001** |
| | Adulthood | -1.880 | 0.167 | 78 | -11.29 | **<0.001** |
| *ife-2* | Intercept (Control) | 4.339 | 0.056 | 2.0 | 76.90 | **<0.001** |
| | Lifelong | 0.045 | 0.054 | 85.3 | 0.83 | 0.411 |
| | Adulthood | 0.125 | 0.054 | 85.1 | 2.32 | **0.023** |

**Table S5. LRS.** The effect of age-specific down-regulation of each gene on lifetime reproductive success. Treatment contrast from generalized mixed-effect models with a Conway-Maxwell-Poisson (CMP) distribution (models with CMP distribution had lowest AIC for all genes, see Table S6).

| Gene | Treatment contrast | coef | SE | d.f. resid. | z | p |
|---|---|---|---|---|---|---|
| *age-1* | Intercept (Control) | 5.580 | 0.033 | 81 | 167.85 | **<0.001** |
| | Lifelong | 0.032 | 0.031 | 81 | 1.04 | 0.299 |
| | Adulthood | 0.034 | 0.031 | 81 | 1.10 | 0.271 |
| *raga-1* | Intercept (Control) | 5.556 | 0.039 | 85 | 141.99 | **<0.001** |
| | Lifelong | -0.053 | 0.031 | 85 | -1.73 | 0.083 |
| | Adulthood | -0.022 | 0.030 | 85 | -0.73 | 0.468 |
| *nuo-6* | Intercept (Control) | 5.482 | 0.082 | 79 | 67.28 | **<0.001** |
| | Lifelong | -1.219 | 0.126 | 79 | -9.66 | **<0.001** |
| | Adulthood | -0.373 | 0.098 | 79 | -3.82 | **<0.001** |
| *ifg-1* | Intercept (Control) | 5.576 | 0.031 | 75 | 181.62 | **<0.001** |
| | Lifelong | -7.650 | 0.599 | 75 | -12.76 | **<0.001** |
| | Adulthood | -2.959 | 0.089 | 75 | -33.41 | **<0.001** |
| *ife-2* | Intercept (Control) | 5.556 | 0.042 | 84 | 133.75 | **<0.001** |
| | Lifelong | 0.028 | 0.033 | 84 | 0.84 | 0.403 |
| | Adulthood | 0.016 | 0.033 | 84 | 0.48 | 0.628 |

**Table S6. Egg size.** The effect of age-specific down-regulation of each gene on egg size. Treatment contrast from mixed-effect models.

| Gene | Treatment contrast | coef | SE | d.f.. | t | p |
|---|---|---|---|---|---|---|
| *age-1* | Intercept (Control) | 1.26 e-3 | 1.07e-4 | 1 | 11.84 | 0.052 |
| | Lifelong | -3.81e-6 | 1.73e-5 | 68.5 | -0.22 | 0.826 |
| | Adulthood | -8.36e-6 | 1.64e-5 | 69.6 | -0.51 | 0.612 |
| *raga-1* | Intercept (Control) | 1.09e-3 | 2.66e-5 | 1.40 | 41.18 | **0.004** |
| | Lifelong | 3.39e-5 | 1.88e-5 | 27.1 | 1.80 | 0.083 |
| | Adulthood | 1.18e-5 | 1.88e-5 | 27.0 | 0.63 | 0.535 |
| *nuo-6* | Intercept (Control) | 1.41e-3 | 6.43e-5 | 1 | 21.89 | **0.022** |
| | Lifelong | 6.11e-5 | 2.45e-5 | 26.2 | 2.50 | **0.019** |
| | Adulthood | -7.27e-5 | 2.44e-5 | 25.8 | -0.30 | 0.768 |
| *ifg-1* | Intercept (Control) | 1.26e-3 | 1.09e-4 | 1 | 11.58 | 0.054 |
| | Lifelong | *No eggs produced* | | | | |
| | Adulthood | 1.95e-4 | 1.59e-5 | 50.3 | 12.31 | **<0.001** |
| *ife-2* | Intercept (Control) | 1.10e-3 | 3.40e-5 | 1.17 | 32.17 | **<0.001** |
| | Lifelong | 1.68e-5 | 1.62e-5 | 30 | 1.03 | 0.309 |
| | Adulthood | 4.00e-5 | 1.59e-5 | 30 | 2.52 | **0.017** |

**Table S7.** Comparisons of models of LRS with different error distribution (Poisson, Conway-Maxwell-Poisson [CMP]), as well as a Poisson model with subject level random effects. The model with the lowest AIC was selected.

| Gene | Distribution | d.f. | dAICc |
|---|---|---|---|
| *age-1* | Poisson | 4 | 111.1 |
| | Poisson + subj. level effect | 5 | 2.0 |
| | CMP | 5 | 0.0 |
| *raga-1* | Poisson | 4 | 105.5 |
| | Poisson + subj. level effect | 5 | 4.0 |
| | CMP | 5 | 0.0 |
| *nuo-6* | Poisson | 4 | 1772.6 |
| | Poisson + subj. level effect | 5 | 97.5 |
| | CMP | 5 | 0.0 |
| *ifg-1* | Poisson | 4 | 53.3 |
| | Poisson + subj. level effect | 5 | 31.0 |
| | CMP | 5 | 0.0 |
| *ife-2* | Poisson | 4 | 161.7 |
| | Poisson + subj. level effect | 5 | 12.8 |
| | CMP | 5 | 0.0 |

**Table S8.** Comparisons of models of Age-specific reproduction with different error distribution (Poisson, Conway-Maxwell-Poisson [CMP]), as well as a Poisson model with subject level random effects. The model with the lowest AIC was selected.

| Gene | Distribution | d.f. | dAICc |
|---|---|---|---|
| *age-1* | Poisson | 11 | 996.4 |
| | Poisson + subj. level effect | 12 | 88.9 |
| | CMP | 12 | 0.0 |
| *raga-1* | Poisson | 11 | 1063.5 |
| | Poisson + subj. level effect | 12 | 122.7 |
| | CMP | 12 | 0.0 |
| *nuo-6* | Poisson | 11 | 601.9 |
| | Poisson + subj. level effect | 12 | 51.6 |
| | CMP | 12 | 0.0 |
| *ifg-1* | *Almost complete cross-separation, not possible to model* | | |
| *ife-2* | Poisson | 11 | 878.7 |
| | Poisson + subj. level effect | 12 | 133.9 |
| | CMP | 12 | 0.0 |
